# Predicting species distributions and community composition using remote sensing products

**DOI:** 10.1101/2020.07.06.185322

**Authors:** Jesús N. Pinto-Ledezma, Jeannine Cavender-Bares

## Abstract

Accurate predictions of species composition and diversity are critical to the development of conservation actions and management strategies. In this paper using oak assemblages distributed across the conterminous United States as study model, we assessed the performance of stacked species distribution models (S-SDMs) and remote sensing products in building the next-generation of biodiversity models. This study represents the first attempt to evaluate the integrated predictions of biodiversity models—including assemblage diversity and composition—obtained by stacking next-generation SDMs. We found three main results. First, environmental predictors derived entirely from remote sensing products represent adequate covariates for biodiversity modeling. Second, applying constraints to assemblage predictions, such as imposing the probability ranking rule, results in more accurate species diversity predictions. Third, independent of the stacking procedure (bS-SDM versus cS-SDM), biodiversity models do not recover the observed species composition with high spatial resolution, i.e., correct species identities at the scale of individual plots. However, they do return reasonable predictions at macroecological scales (1 km). Our results provide insights for the prediction of assemblage diversity and composition at different spatial scales. An important task for future studies is to evaluate the reliability of combining S-SDMs with direct detection of species using image spectroscopy to build a new generation of biodiversity models to accurately predict and monitor ecological assemblages through time and space.

## 1. Introduction

Species diversity and composition vary in space and time as a consequence of historical biogeographic and environmental factors and ongoing ecological processes. Since the last millennium, however, rising human population and activities have been major drivers of environmental change on Earth (Dale et al., 2000; Lewis and Maslin, 2015; Pinto-Ledezma and Rivero Mamani, 2014; Tilman et al., 2017). As a consequence, biodiversity is changing at a pace solely compared to the major extinction events recorded in the geologic history (Allen et al. 2020; IPBES, 2019; Lewis and Maslin, 2015; Pimm et al., 2006). These rapid changes in biodiversity are impacting the capacity of ecosystems to provide services to humanity that ultimately influence our well-being (Cavender-Bares et al., 2015a; Chaplin-Kramer et al., 2019; Díaz et al., 2019; Pecl et al., 2017). Thus, detecting and monitoring species diversity and composition is critical to developing effective management strategies and conservation actions facing global change (Jetz et al., 2019; Pinto-Ledezma and Rivero Mamani, 2014; Purvis et al., 2018; Watson et al., 2020) that move us towards international biodiversity goals, including those posed by the Intergovernmental Science-Policy Platform on Biodiversity and Ecosystem (IPBES) and the parallel United Nations’ (UN) Sustainable Development Goals (UN-SDGs) (Díaz et al., 2019) and the upcoming post-2020 Convention on Biodiversity goals.

Different approaches have been proposed for assessing and monitoring the spatial and temporal patterns of species diversity and distribution (Jetz et al., 2019; Mateo et al., 2017; Pereira et al., 2013), including biodiversity models—i.e., models that aim to predict species diversity and composition (Ferrier and Guisan, 2006; Guisan and Rahbek, 2011) and remote sensing technologies and products (Caughlin et al., 2016; Cavender-Bares et al., 2020; Fawcett et al., 2018; Féret and Asner, 2014; Randin et al., 2020; Turner, 2014). Among biodiversity models, the approach of Stacked Species Distribution Models (S-SDMs) has been successfully implemented for predicting species diversity and composition (D’Amen et al., 2015; Pottier et al., 2013; Zurell et al., 2020). In building S-SDMs, individual SDM are modeled as a function of environmental predictors—usually derived from interpolated climate surfaces. Subsequently, these models are stacked to produce species diversity and composition assemblage predictions (Ferrier and Guisan, 2006; Pottier et al., 2013). However, despite successful S-SDMs predictions, most previous studies were restricted to small areas (D’Amen et al., 2018, 2015; Pottier et al., 2013), limiting the potential of these models for biodiversity prediction and monitoring over large geographic areas, including at continental scales. Moreover, the uncertainty associated with the underlying data (i.e., occurrence records and environmental covariates) can negatively impact species model predictions (Dobrowski et al., 2011; Soria-Auza et al., 2010) and consequently the reliability of these kinds of biodiversity models.

Remote sensing products (RS-products) have been increasingly used to derive metrics that allow tracking biodiversity from space (Lausch et al., 2018; Rocchini et al., 2016; Schulte to Bühne and Pettorelli, 2018; Turner, 2014), monitoring the state of human impacts (Hansen et al., 2013; Pinto-Ledezma and Rivero Mamani, 2014), as predictors for describing large patterns of species diversity (Hobi et al., 2017; Radeloff et al., 2019) or to derive Essential Biodiversity Variables, i.e., measures that allow the detection and quantification of biodiversity changes (Fernández et al., 2020; Pereira et al., 2013). Despite their high spatial and temporal resolution, quasi-global coverage and range of data products (e.g., precipitation, plant productivity, biophysical variables, land cover), RS-products have been rarely used as predictors for biodiversity models (Pinto-Ledezma and Cavender-Bares, 2020; Randin et al., 2020; Saatchi et al., 2008). RS-products have been dubbed important “next-generation” environmental predictors in biodiversity models (He et al., 2015), given that remote sensing continuously captures an increasing range of Earth’s biophysical features at global scale (Cord et al., 2013; Guanter et al., 2014; Jetz et al., 2016), avoiding the uncertainty associated with environmental predictors derived from traditional climatic data. Furthermore, species models derived from RS-products perform as well as from those derived from interpolated climate surfaces have the potential to provide predictions with greater spatial resolution (Pinto-Ledezma and Cavender-Bares, 2020).

A crucial challenge for detecting and monitoring biodiversity is the integration of modeling approaches with RS-products in order to build next-generation biodiversity models at different spatial and temporal scale and levels of organization (He et al., 2015; Randin et al., 2020). Therefore, the general aim of this paper is twofold: (1) to combine S-SDMs and RS-products in order to build next-generation biodiversity models, and (2) to perform a rigorous evaluation of biodiversity model predictions. In doing so, we applied a sequential procedure that allows the prediction and evaluation of species assemblage diversity and composition at continental scale (Fig. 1). This procedure was implemented using the oak clade (genus *Quercus*) as a model study within the geographical area corresponding to the conterminous United States. We focused on this clade because oaks are widely distributed forest dominants, and they have the highest species richness and highest total biomass of any woody group in the conterminous U.S. (Cavender-Bares, 2019). Nevertheless, individual species have contrasting environmental distributions and range sizes. These attributes make the oaks both an important study system and a useful one for testing next generation SDMs.

**Fig. 1.**
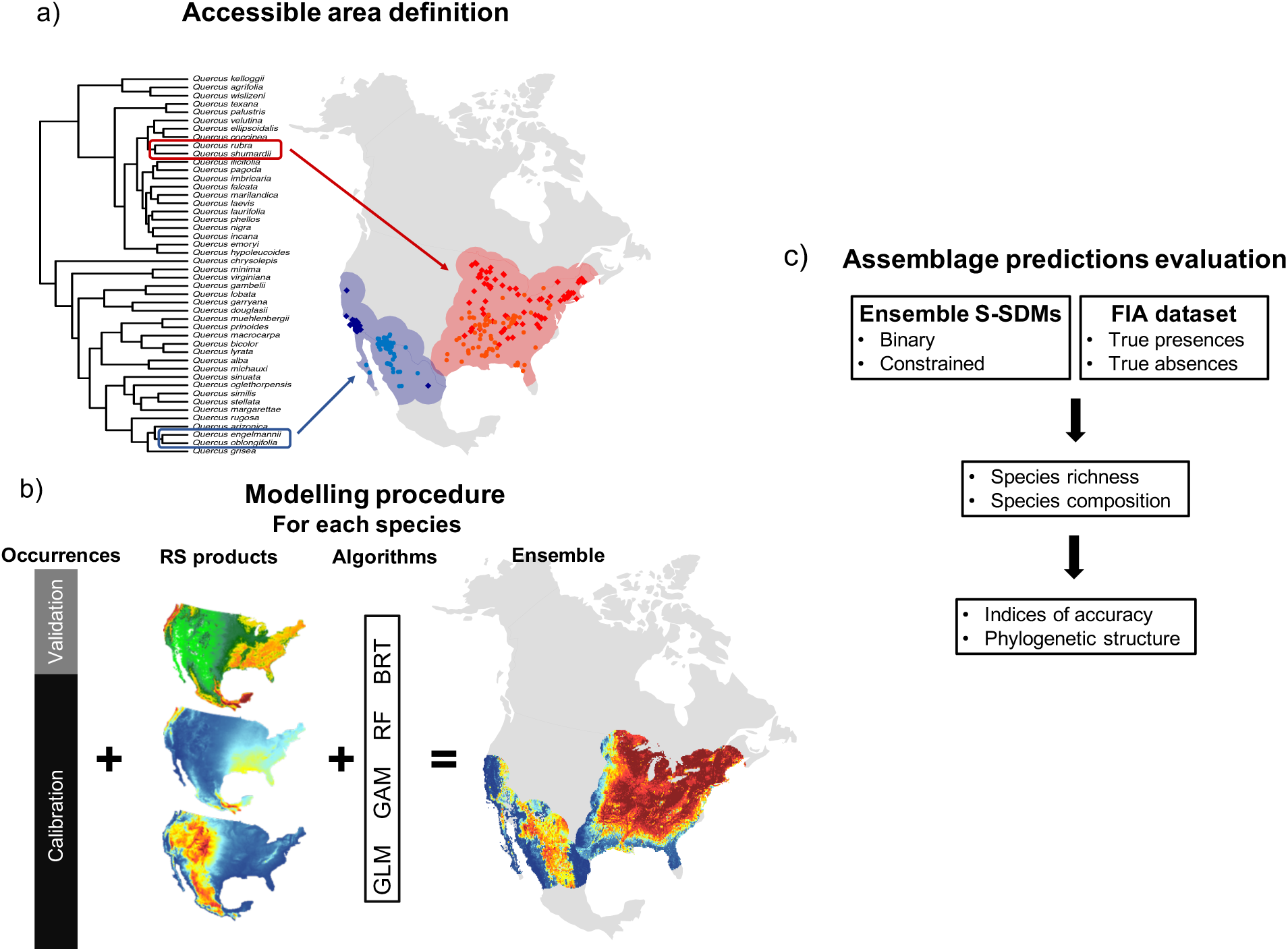
Implementation of the sequential procedure to construct and evaluate biodiversity model predictions. The first step defines the accessible area for each oak species. In this study, the accessible area is defined as the overlap of occurrences (confirmed presences) of sister species (a). The second step involves implementing an ensemble modeling approach for individual oak species but constrained to the accessible area established in the step one (b). Lastly, individual model predictions are stacked (S-SDMs) to obtain species richness, phylogenetic structure and composition predictions. These predictions are then evaluated using an independent dataset of species diversity and composition obtained from FIA (c).

## 2. Methods

### 2.1. Species occurrence dataset

This genus *Quercus* includes 91 species widely distributed across U.S. and showing a marked longitudinal species diversity gradient, with high species concentration in south-eastern North America. Our main occurrence dataset was assembled in a previous study (Hipp et al., 2018). We completed the main dataset using the Integrated Digitized Biocollections (iDigBio, data downloaded between 15 and 18 April, 2019) and collections from the second author (JCB) (Cavender-Bares et al., 2015b). All occurrence data were visually examined and any localities that were outside the known geographical range of the species, in unrealistic locations (e.g., water bodies, crop fields) or in botanical gardens were discarded for accuracy. In addition, to avoid problems of spatial sampling bias and spatial autocorrelation we thinned the occurrence records of each species using a spatial thinning algorithm with thinning distance of 5 km (Aiello-Lammens et al., 2015). The final dataset included 132 ± 211 (SD) unique presence data per species and 77.099% of the species had more than 25 records and 22.901% at least 10 presence records.

### 2.2. Environmental data derived from Remote sensing

Remote sensing products used as environmental inputs to our models included Leaf Area Index (LAI), obtained from MODIS Terra/Aqua MOD15A2 over a 15-year period (2001 – 2015) using the interface EOSDIS Earthdata (Myneni et al., 2002); precipitation from Climate Hazards group Infrared Precipitation with Stations (CHIRPS); and altitude or mean elevation from the Shuttle Radar Topography Mission (SRTM). The 15-year LAI composite provides a representation of the spatial variation of biophysical variables of different components of vegetation and ecosystems over the course of this time frame (Hobi et al., 2017; Pinto-Ledezma and Cavender-Bares, 2020). LAI strongly co-varies with the physical environment; for instance, higher LAI is associated with warmer, wetter and stable environments, whereas lower LAI with cooler, drier and less stable environments (Gower et al., 1999; Myneni et al., 2002; Reich, 2012). In particular, LAI has a strong relationship with climate and captures the dynamics of the growing season, including vegetation seasonality—important for characterizing plant species geographical ranges (Cord et al., 2017). We posit, therefore, that the spatial and temporal variation in LAI—half of the total green leaf area per unit of horizontal ground surface area (Xiao et al., 2014)—represents an important variable for determining species distributions that integrates across biotic and abiotic factors (Peterson, 2011; Soberón, 2007). CHIRPS is a quasi-global, high resolution and daily precipitation dataset designed for monitoring drought and global environmental land change (Funk et al., 2015). SRTM is a high-resolution digital elevation model of the Earth that has been used for mapping and monitoring the earth’s surface (Farr et al., 2007) and an important variable for predicting species distributions and community composition (Dubuis et al., 2011; Pottier et al., 2013).

### 2.3. Species and community composition validation data

Ecological niche models represent abstractions of the environmentally suitable areas for species to maintain long-term viable populations (Peterson, 2011). The presence of a species in a locality or a grid cell informs us about the areas that are environmentally suitable for that species (Lobo et al., 2010; Peterson, 2011). Its absences inform us of the opposite pattern—namely those areas that are either not environmentally suitable for the species or are the result of historical contingences, biotic interactions that constraint the species presence even if the physical environment is suitable, recent extirpation events such as those caused by land use change, or simply because the species was not detected (Lobo et al., 2010; Peterson, 2011; Václavík and Meentemeyer, 2009). Here, using an independent dataset of true presences and absences from the Forest Inventory and Analysis (FIA) program of the U.S. Forest Service (Smith, 2002), we evaluate the predictions of individual oak species models and also the community composition predictions. A standard FIA plot is designed to cover an area of 6052 m^2^ (∼0.6 ha); within each plot, four circular subplots of 168.3 m^2^ were established with one center subplot and three equidistant subplots. Within each subplot every tree species with stem >2.5 cm diameter at breast height is recorded. Although the surveys are intensive, not all trees in the plot are measured because the four subplots do not cover the total area of each surveyed plot (https://www.fia.fs.fed.us/library/database-documentation/current/ver70/FIADB%20User%20Guide%20P2_7-0_ntc.final.pdf). Data of this type is commonly used in community ecology, community phylogenetics and biogeography (e.g., Cavender-Bares et al., 2018; Potter et al., 2019).

Oak species from the FIA dataset was obtained from a previous study (Cavender-Bares et al. 2018). FIA is a continuous forest census that aims to determine the extent, condition, volume, growth, and use of trees in the U.S. Shrub species are largely excluded from the FIA inventory. This dataset includes a total of 112,183 plots of 6,052 m^2^ and 49 of the 91 oak species found in naturally assembled forests (Cavender-Bares et al., 2018). From this data set we filtered assemblages to retain only those with at least three species. The final FIA dataset used in this study includes a total of 13,263 plots and 49 oak species in the conterminous U.S.

### 2.4. Modeling framework

To obtain reliable SDM predictions it is necessary to define the accessible area for a species in geographical space, i.e., the area that has been accessible to the species within a given period of time (Barve et al., 2011; Peterson, 2011). We defined the accessible area for each oak species as the combined area of the geographical ranges of sister species or the overlap of geographical ranges of a given species and its phylogenetically closest relatives (Fig. 1a). Note that the geographical ranges were defined using the occurrence records of each oak species. To do so, we first identified sister species using the published phylogeny of American oaks (Hipp et al., 2018). This phylogeny was constructed using genome-wide restriction site-association DNA sequencing (RAD-seq) and is the most highly resolved phylogenetic hypothesis for the clade in North America (Hipp et al., 2018). The rationale behind this approach is to define a large area that a species can colonize based on species environmental tolerances—i.e., under the assumption that oak species share environmental tolerances (Hipp et al., 2018; Cavender-Bares et al., 2018). Importantly, this procedure provides a conservative spatial representation of the environmental space in which a given species have potentially dispersed and detected.

Prior to model calibration, we masked the environmental variables to the extent of the defined species accessible area. This procedure was repeated for each oak species. Using an ensemble framework (Araujo and New, 2007; Diniz-Filho et al., 2009) and following Pinto-Ledezma and Cavender-Bares (2020), we modeled the ecological niche for each oak species (Fig. 1b). Under this framework, the predicted species ecological niche or distributional area is the outcome of combining the results of different modeling algorithms (Araujo and New, 2007; Diniz-Filho et al., 2009). Within this framework we fit four species models including two statistical methods (generalized linear models [GLM] and generalized additive models [GAM]) and two machine learning methods (Boosted regression trees [BRT] and Random Forest [RF]). Model evaluation was performed using a K-fold cross-validation (James et al., 2013). This approach randomly splits the data into K-folds or K-groups of relatively equal size and then fits every model K times. At each step (K-folds) K-1 folds are used for training and remaining 1-fold for testing (James et al., 2013). We used 5 folds to split our data, i.e., 80% and 20% of the data were used for testing and training, respectively. The final ensemble model prediction for each species (Fig. 1b) was obtained using weighted averaging based on true skill statistic (Allouche et al., 2006). The ensemble procedure including model calibration, evaluation and ensemble prediction was performed using the R package sdm (Naimi and Araújo, 2016).

To obtain assemblage composition and species richness predictions we applied two stacking species distribution models (S-SDM) procedures (i.e., binary and constrained binary) (Fig. 1c). These procedures return the predicted species composition and richness within each assemblage (i.e., grid cell) across a geographical domain (Ferrier and Guisan, 2006; Guisan and Rahbek, 2011). More specifically, binary S-SDM (bS-SDM) predictions were obtained by converting individual species ensembles into binary predictions or occurrence probabilities by using its corresponding TSS-maximization threshold (Scherrer et al., 2018; Zurell et al., 2020). Constrained binary S-SDM (cS-SDM) predictions were obtained by constraining each assemblage applying a probability ranking rule (PRR). PRR emulates ecological assembly rules by ranking the species in each assemblage based on the occurrence probability obtained from each species and the number of species per assemblage. The species with the highest probabilities in an assemblage is selected until the number of species in an assemblage, based on observed data, is reached (D’Amen et al., 2015; Scherrer et al., 2018). We used the maximum number of species per assemblage from the FIA dataset as assemblage-level constraint for cS-SDM estimations.

### 2.5. Evaluating species and assemblage predictions

Given that the FIA dataset used in this study includes only 49 out of the 91 oak species in the U.S., we restricted the species and assemblage prediction evaluations to those 49 species (Supplementary table 1). The performance of species predictions was assessed using the Boyce index, as implemented in Hirzel et al. (2006). This index varies from -1 to +1, in which negative values indicate counter predictions and positive values indicate that model prediction are consistent with the observed distribution of presences, while values close to zero suggest that model predictions are not different from random (Boyce et al., 2002; Hirzel et al., 2006).

To evaluate the performance of species assemblage predictions, we used four different metrics that are commonly used for this purpose (Pottier et al., 2013; Scherrer et al., 2018; Zurell et al., 2020). The metrics used were: a) the deviation of the predicted species richness to the observed (SR deviation); b) the proportion of correctly predicted as present or absent (Prediction success); c) true skill statistic (TSS) and d) similarity between the observed and predicted community composition (Sørensen index). Assemblage metric evaluations were performed using the R package ecospat (Di Cola et al., 2017) using the matrices of the two assemblage predictions, i.e., binary and PRR (Fig. 1c) and 1000 iterations. Note that the SR deviation was estimated only for the bS-SDM predictions given that we used the FIA dataset in constraining the number of species per assemblage in the cS-SDM construction.

We also investigated the phylogenetic structure of oak assemblages for both the FIA dataset and the predicted assemblages, in order to explore the performance of predicted oak assemblages in recovering similar patterns of phylogenetic structure as the observed assemblages. We defined phylogenetic structure as the mean phylogenetic distance (MPD) (Webb et al., 2002). To facilitate comparison between the two datasets, we summarized the results using standardized effects sizes (SES), which compare the observed value of an assemblage (MPD) to the mean expected value under a null model, correcting for their standard deviation. SES values >0 and <0 indicate phylogenetic clustering and overdispersion, respectively (as in Webb et al., 2002). In SES calculations we randomized the tips of the phylogeny to generate random communities (taxon shuffle null model). All phylogenetic structure calculations were conducted using customized scripts and core functions from the PhyloMeasures (Tsirogiannis and Sandel, 2016) package in R.

Given the mismatch between the grain size of the observed assemblages (6,052 m^2^ or ∼0.006 km) and the grain size of the predictions (1 km), we were also interested in exploring the effect of increasing grain size in assemblage predictions. Accordingly, we aggregated the observed FIA assemblages into larger assemblages, ranging from 30 arcseconds (∼1 km), 2.5 degrees (∼25 km) and 0.5 degrees (∼55 km), and repeated all the assemblage level analyses described above. Data aggregation was performed using the R package letsR (Vilela and Villalobos, 2015).

### 2.6. Statistical analysis

Using a Bayesian counterpart of Pearson’s correlation test, we evaluated the relationship between species richness and phylogenetic assemblage structure obtained from both FIA and predicted datasets. We chose this Bayesian alternative because it allow robust parameter estimations and account for outliers in the data. We also used Bayesian-ANOVAs to test for differences in species assemblage predictions between different modelling procedures (i.e., stacking procedures), using plots as a random variable to correct for potential repeated measures. Using Maximum A Posteriori (MAP) p-values (Mills and Parent, 2014), we then evaluated the evidence for those differences. Note that all analyses were performed for each grain separately. Bayesian-ANOVAs was performed using the R package rstanarm (Goodrich et al., 2020), and the Bayesian Pearson’s correlation test was implemented using the R package BayesianFirstAid (https://github.com/rasmusab/bayesian_first_aid). All analyses were performed using 4 sampling chains for 10,000 generations, discarding 20% of each run as burnins. MAP-based p-values was estimated as implemented in the R package bayestestR (Makowski et al., 2019).

## 3. Results

### 3.1. Species predictions

Oak species predictions were successfully calibrated for all species (Fig. S1), showing mid to high prediction accuracy as measured by the AUC (area under the receiver operating characteristic curve) and the True skill statistic (TSS) metrics (Ensemble mean ± SD: AUC 0.858 ± 0.056, TSS 0.703 ± 0.049; Fig. S1). The performance of species predictions also showed high accuracy: all species presence values were greater than 0.5 as measured by the Boyce index (mean ± SD: 0.897 ± 0.140; Table S1), indicating that ensemble model predictions are consistent with the independent observed presences from the FIA dataset.

### 3.2. Assemblage predictions

Mean observed richness per assemblage at the observed grain size (i.e., FIA plots of 6052 m^2^) was 3.430 (SD = 0.922, max = 10, min = 3), and mean richness prediction per assemblage was 6.104 (SD = 2.443, max = 12, min = 1). Mean observed richness at 30 arc seconds (i.e., spatial resolution of the predictors) was 4.039 (SD = 1.299, max = 12, min = 3), while the prediction for richness was 8.322 (SD = 3.474, max = 16, min = 3). Bayesian correlations revealed low but positive correlations between the observed and the predicted richness derived from the bS-SDM at the observed grain size (ρ = 0.26 [0.24:0.27, 95% confidence interval (CI)], Fig. 2a). Interestingly, the relationship between the observed and predicted richness increased dramatically as the grain size increased (Fig. 2b-d).

**Fig. 2.**
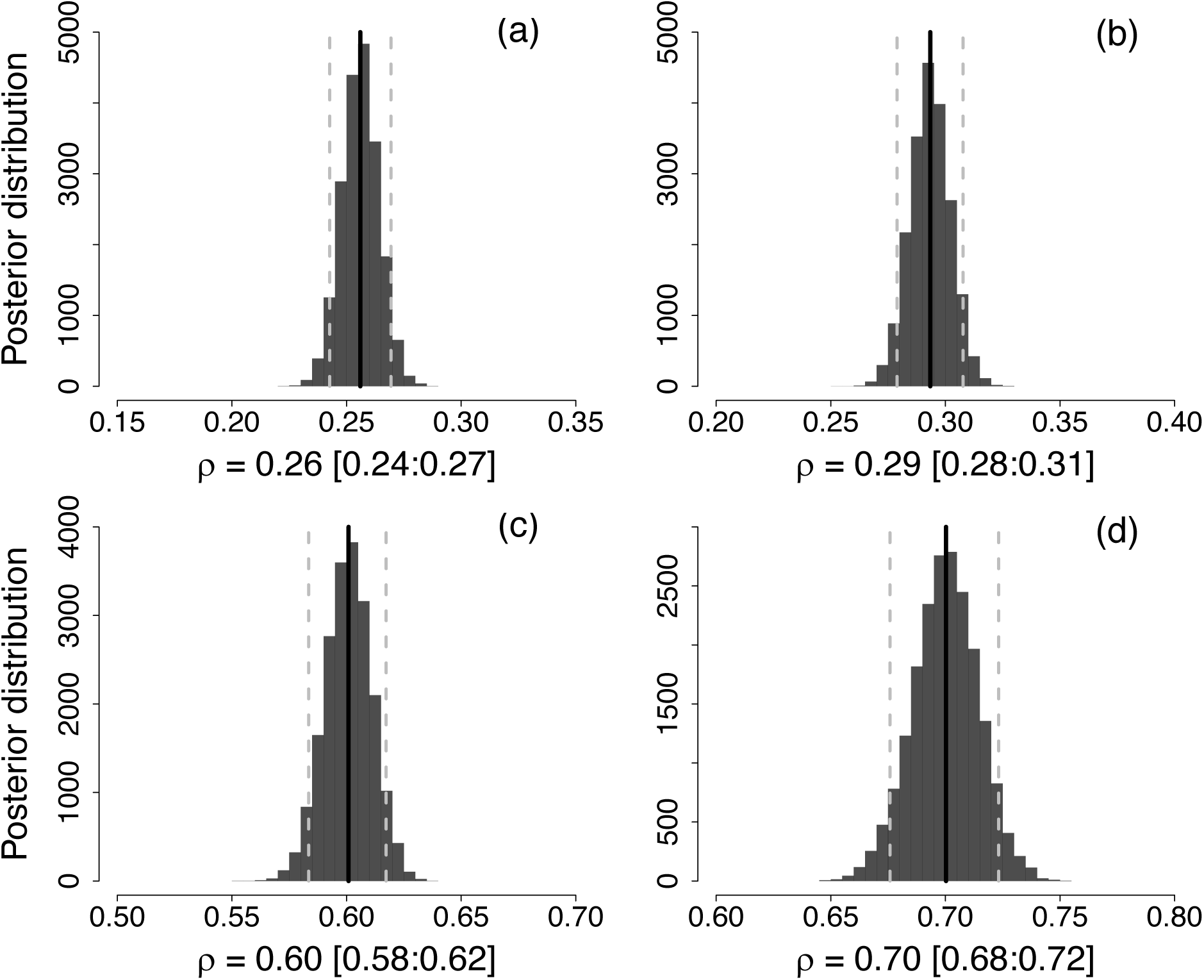
Bayesian Pearson’s correlation coefficients between the predicted and observed species richness at different grain size. Observed grain size or FIA plots of ∼0.006 km^2^ (a), 30 arcseconds or 1 km^2^ aggregation (b), 0.25 degrees or ∼25 km^2^ aggregation(c), and 0.5 degrees or ∼55 km^2^ aggregation (d). Posterior distribution show ρ values from 4 sampling chains and 10,000 generations. The vertical black line represent the median (and 95% confidence interval, vertical gray dotted lines) ρ value from posterior distribution.

Likewise, mean absolute deviation between observed species richness and species richness predictions derived from the bS-SDM was 0.246 (SD = 0.204, max = 0.939, min = 0.000) and decreased as grain size increased (Fig. 3a). The mean assemblage prediction success for the bS-SDM was 0.885 (SD = 0.047, max = 0.983, min = 0.744; Fig. 3b) and for the cS-SDM 0.941 (SD = 0.037, max = 1, min = 0.739; Fig. 3b). Average assemblage TSS over all localities for bS-SDM were 0.436 (SD = 0.140, max = 1, min = 0.021; Fig. 3c) and 0.562 (SD = 0.276, max = 1, min = 0.121; Fig. 3c) for cS-SDM. Mean assemblage similarity estimated using the Sørensen index was 0.551 (SD = 0.125, max = 0.905, min = 0.1; Fig. 3d) for bS-SDM and 0.594 (SD = 0.258, Max = 1, min = 0.01) for cS-SDM. As expected, the accuracy of assemblage predictions increased with increasing grain size (Fig. 3). In addition, although the mean values of the metrics used to evaluate the accuracy in assemblage predictions were relatively similar between the two stacking procedures (bS-SDM vs cS-SDM), Bayesian-ANOVAs revealed strong evidence in favor for cS-SDM in predicting assemblage composition, in other words, cS-SDM stacking procedure outperformed bS-SDM. The higher performance of the cS-SDM procedure is consistent across all grain sizes, except for TSS at the grain size 0.5 (Fig. 3c), which can be considered anecdotal.

**Fig. 3.**
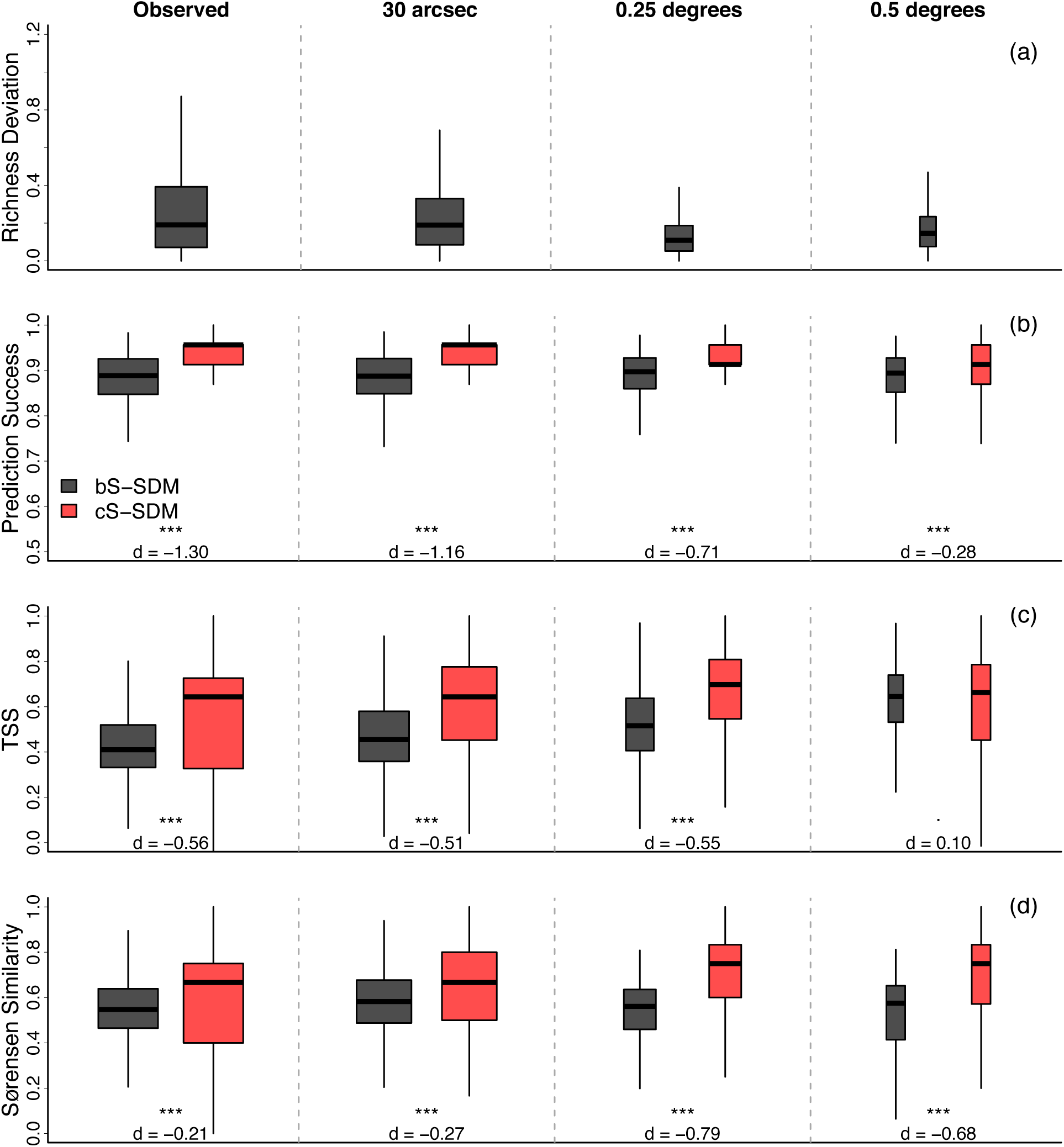
Accuracy of assemblage composition predictions at different grain size. (a) Richness deviation, measured as the absolute deviation in assemblage richness predictions divided by the maximum observed FIA plot species richness, (b) prediction success or accuracy in assemblage predictions, (c) True skill statistics, TSS, and (d) Sørensen similarity. Significant differences between stacking procedures (bS-SDM versus cS-SDM) were evaluated using Bayesian ANOVAs with FIA plots as random effect (significance levels for the Maximum A Posteriori p-values are: ***p < 0.001, **p < 0.01, *p < 0.05, •p < 0.1). We also show Cohen’s *d* effect size for each comparison. Note that the width of each boxplot is weighted by the sample size. Boxplots are ordered according grain size, from the observed grain size or FIA plot of ∼0.006 km^2^ (left-hand boxplots) to the largest grain size (right-hand boxplots), i.e., FIA plots aggregated at 0.5 degrees or ∼55 km^2^.

We were also interested in evaluating whether species composition of the assemblage predictions returned similar patterns of phylogenetic structure as the observed assemblages. Accordingly, the average SES-MPD at the observed grain size for the FIA dataset was -0.471 (SD = 1.130), for the bS-SDM -1.915 (SD = 1.396) and -0.201 (SD = 0.906) for the cS-SDM. In addition, most of the predicted assemblages presented negative SES-MPD values (Fig. 4), indicating that the dominant pattern of phylogenetic structure in the predicted oak assemblages is overdispersion, a pattern that is observed in natural oak assemblages. Bayesian correlations showed low but positive associations between the phylogenetic structure obtained from the observed and both assemblage predictions (ρ = 0.15 [0.14:0.17] and ρ = 0.26 [0.24:0.27], median rho and 95% CI estimated from FIA vs bS-SDM and FIA vs cS-SDM, respectively) at the observed grain size (Fig. 5a-b). Similar to the richness predictions, the correlation between the phylogenetic structure among datasets increased sharply with increasing grain size (Fig. 5d-l).

**Fig. 4.**
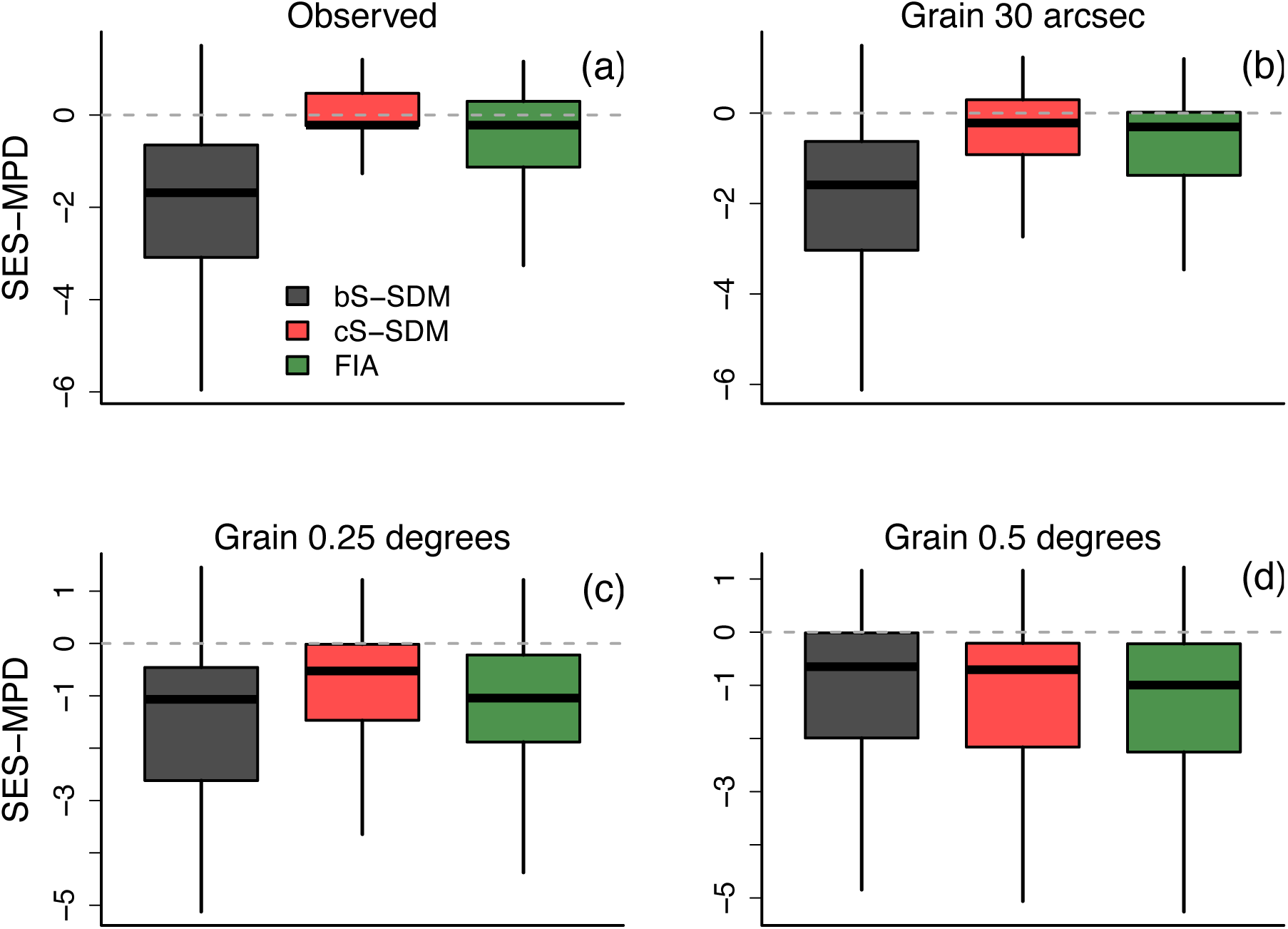
Phylogenetic structure (standardized effect size–mean phylogenetic distance [SES-MPD]) of natural and predicted oak assemblages at four grain sizes. SES-MPD at observed grain size (∼0.006 km^2^) (a), SES-MPD at 30 arcseconds or 1 km^2^ aggregation (b), SES-MPD at 0.25 degrees or ∼25 km^2^ aggregation(c), and SES-MPD at 0.5 degrees or ∼55 km^2^ aggregation (d).

**Fig. 5.**
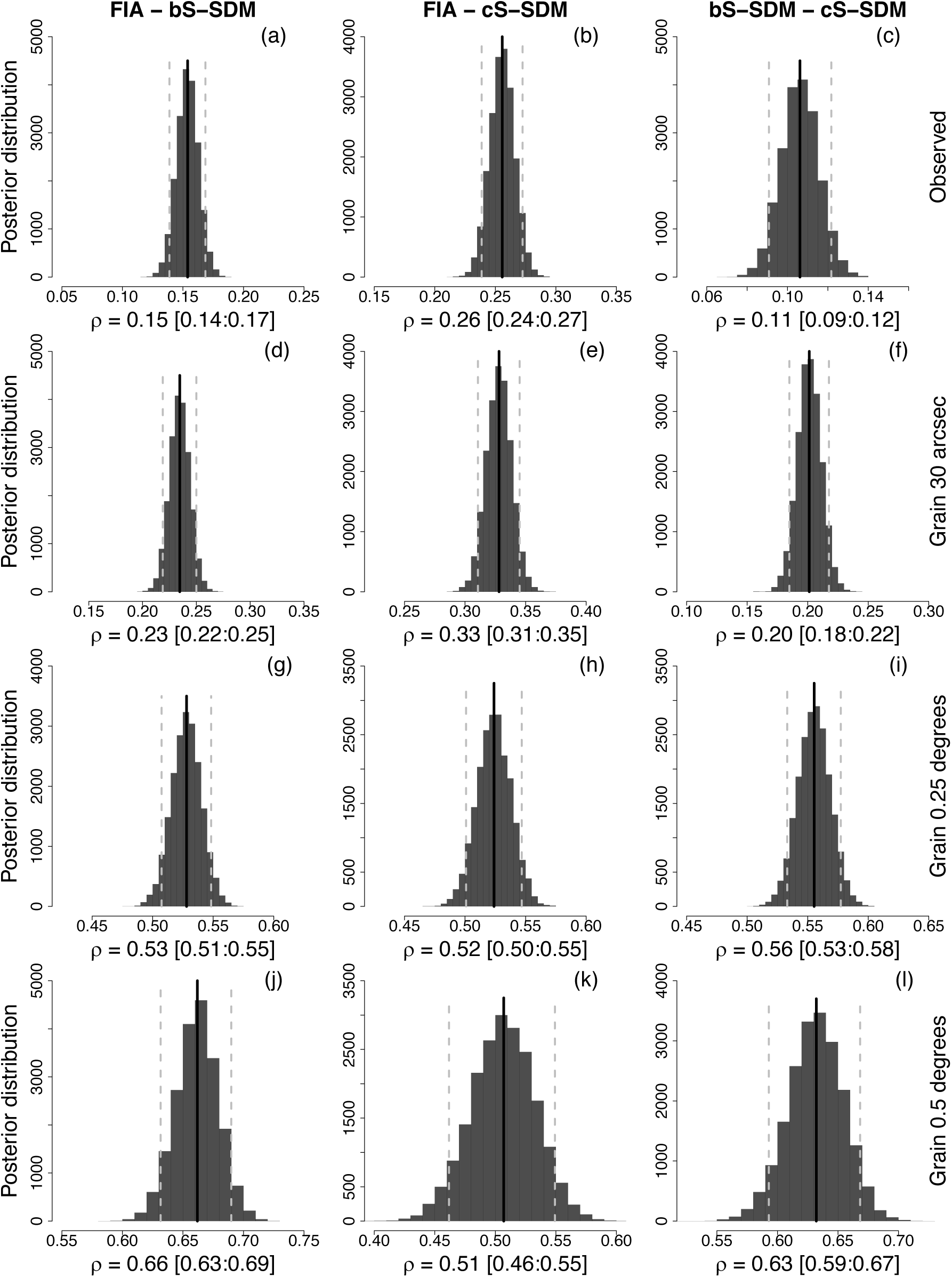
Bayesian Pearson’s correlation coefficients between the predicted (bS-SDM and cS-SDM) and observed assemblage phylogenetic structure at different grain size. (a-c) Observed grain size or FIA plots of ∼0.006 km^2^, (d-f) 30 arcseconds or 1 km^2^ aggregation, (g-i) 0.25 degrees or ∼25 km^2^ aggregation, and (j-l) 0.5 degrees or ∼55 km^2^ aggregation. Posterior distributions show ρ values from 4 sampling chains and 10000 generations. The vertical black line represent the median (and 95% confidence interval, vertical gray dotted lines) ρ value from posterior distribution.

## 4. Discussion

The use of RS-products in biodiversity models has been hailed as a transformative approach for providing simple and flexible biodiversity predictions (He et al., 2015; Randin et al. 2020). In this study we evaluated the reliability of S-SDM—a biodiversity model (Ferrier and Guisan, 2006)—based on individual SDMs derived from RS-products only, to predict biodiversity in terms of species richness and composition within local assemblages. Our results indicate that the combination of S-SDM and RS-products perform well in predicting plot-level biodiversity as assessed by several important metrics. However, when we compared the S-SDM predictions of phylogenetic structure of oak assemblages at the plot-level with the observed structure (measured in terms of species richness and mean phylogenetic distance), the predictions tended to overestimate the diversity, even though the predicted and observed metrics were positively correlated. This result is consistent with the stacking procedure. We also found that when oak assemblages were aggregated at large grain sizes, S-SDM recovered the observed pattern well, i.e., patterns of predicted diversity were highly similar to those obtained from the aggregated FIA plots (Fig. 2c-d, 5g-l). Consequently, our results show that biodiversity models in combination with RS-products do not necessarily predict biodiversity well at the plot-level scale, but they do well at larger grain sizes. In other words, the S-SDMs that use RS-products as environmental predictors are very useful for detecting macroecological patterns of biodiversity. These results spur new insights for biodiversity prediction and monitoring at different spatial scales.

Biodiversity models, such as the S-SDMs, depend on the reliability of individual species models (SDM) (Guisan and Rahbek, 2011). SDMs represent abstractions of species Hutchinsonian niches, i.e., species distributions are constrained by both, the abiotic environment and biotic interactions with other species (Hutchinson’s duality *sensu* Colwell and Rangel, 2009; see also, Peterson et al., 2011; Pinto-Ledezma and Cavender-Bares, 2020). Our results show that individual oak SDMs constructed using environmental covariates derived from RS-products, have good accuracy (Fig. S1). However, by stacking a suite of SDMs to obtain assemblage composition predictions, no biotic constraints are considered. Consequently, S-SDM predictions tend to overestimate the number of species within local assemblages (Guisan and Rahbek, 2011; Mateo et al., 2017). We found that species richness predictions using S-SDM overpredict the number of oak species that are observed in naturally assembled oak communities (Fig. 3a). This might be related with the proportion of common (high prevalence) and rare (low prevalence) species that are co-occurring in these assemblages (Fig. S2), that in turn is affected by microhabitat variables that allow oak species to differentiate locally. Indeed, soil moisture (measured in 20×50m plots on the ground) explain oak species trait distributions better than climate variables (Cavender-Bares et al. 2020).

In addition, it has been suggested that incorporating some ecological assembly rules, e.g., the probability ranking rule approach (PRR), in S-SDM predictions can minimize assemblage overpredictions (Calabrese et al., 2014; D’Amen et al., 2015; Guisan and Rahbek, 2011). Our results corroborate this assumption; specifically, predicting species assemblages by simply overlaying individual SDMs, tends to overpredict the number of species present in local assemblages, especially at finer grains (Fig. 2a-b, 3a). Moreover, through the implementation of PRR in S-SDM predictions—i.e., by constraining the number of species present within local assemblages (Guisan and Rahbek, 2011)—we found that cS-SDM outperform bS-SDM assemblage predictions across all metrics used and grain sizes evaluated (Fig. 3b-d).

While applying constraints to assemblage predictions may be advantageous for predicting species richness (Guisan and Rahbek, 2011; Mateo et al., 2017), this is not the case for predicting species identities or composition within assemblages, particularly at finer grains. Indeed, we found that neither bS-SDM and cS-SDM accurately recover species identities at the grain size of plots (Fig. 5a-b) or at the grain size of the predictors variables (Fig. 5d-e). Not surprisingly, predictions of assemblage richness and composition improved dramatically with increasing grain size (Fig. 2c-d and 5g-k), a pattern observed independently of the implementation of PRR (cS-SDM, Fig. 5h and 5k) or its non-implementation (bS-SDM, Fig. 5g and 5j). These contrasting results might be a consequence of the fact that S-SDMs were primarily designed to assess macroecological patterns of species diversity (Ferrier and Guisan, 2006; Mateo et al., 2017). In addition, although RS-products are useful covariates for spatial modelling of biodiversity, they also reduce the uncertainty associated with environmental covariates in model predictions (Pinto-Ledezma and Cavender-Bares, 2020); consequently, the coarse grain of current RS-products hampers accurate predictions of biodiversity at ecological scales (Gamon et al., 2020; Jetz et al., 2016, Pinto-Ledezma and Cavender-Bares, 2020), in other words, the broad spatial resolution of current RS-products (usually with a pixel size of 1 km) limits our ability to capture environmental features at ecological scales (e.g., microtopography, soil moisture, landscape structure) that ultimately determine the coexistence of species locally. Furthermore, we acknowledge that we evaluated only one type of biodiversity model—S-SDM using two stacking procedure—and that other biodiversity models such as the Joint-SDM (Pollock et al., 2014), could potentially improve biodiversity predictions. Nevertheless, a recent study suggests that Joint-SDM do not improve species assemblage predictions (Zurell et al., 2020). Further research is needed to correctly predict biodiversity at the assemblage level.

The evaluation presented here is meant to stimulate further methodological and empirical research to better predict biodiversity at different spatial and temporal scales and levels of organization. A promising approach for this purpose is the hierarchical modeling of species distributions (Mateo et al., 2019; Petitpierre et al., 2016). H-SDMs allow the simultaneous modeling spatial patterns of biodiversity at ecological and regional scales. In constructing HSDMs, individual SDMs are first fit at regional and landscape scales, and the two predictions are fused to obtain a single species SDM, i.e., the regional model is rescaled to the landscape model (Mateo et al., 2019; Petitpierre et al., 2016). This is an interesting approach because it fuses the benefits of macroecological covariates (derived from climatic surfaces or RS-products) with those derived from high-resolution remote sensing products (Mateo et al., 2019).

An unparalleled alternative is the direct detection of species using imaging spectroscopy (Cavender-Bares et al., 2017; Schweiger et al., 2020). For example, leaf-spectra variation among individual plants obtained using imaging spectroscopy provide sufficient information for the correct assignment of populations to species to clades (Cavender-Bares et al., 2016; Meireles et al., 2020; Schweiger et al., 2020) and airborne imagery accurately assigns vegetation canopies to species (Foster and Townsend, 2004). Current and forthcoming hyperspectral images from DESIS sensors and forthcoming SBG and CHIMES sensors, among others (Alonso et al., 2019; Stavros et al., 2017; Turner, 2014) will capture information from the Earth at fine spectral resolution, allowing the estimation of plant traits, plant nutrient content, biophysical variables (e.g., leaf area index, biomass), that can be used for direct detection of functional and perhaps community diversity from space (Jetz et al., 2016). Combining the detection of species using imaging spectroscopy with SDMs to build a new generation of biodiversity models, may open new avenues for the accurately assignment of species and assessment of ecological assemblages through space and time (Féret and Asner, 2014; Meireles et al., 2020; Randin et al., 2020), a critical hurdle to overcome in addressing the challenges posed by the global change.

## 5. Conclusion

Recent review papers (He et al., 2015; Randin et al., 2020) proposed the applicability of remote sensing data as environmental covariates in the construction of next-generation SDMs. Here using environmental covariates derived completely from remote sensing data, we modelled the distribution of oaks (genus *Quercus*) and predicted the number and composition of species within assemblages at different grain sizes. Despite high variability in the predictions, modeled oak assemblages showed phylogenetic overdispersion, indicating that models recovered the observed pattern of distantly related oak species co-occurring more often than expected. Overall we conclude that species richness can be predicted with high accuracy by applying constraints to the predictions, however, accurate predictions of species identities is still an evolving task. We suggest two alternatives (i.e., H-SDMs and the direct detection of species using imaging spectroscopy) that might increase the accuracy in assemblage composition predictions, hence, future studies should evaluate the reliability of these two alternatives using different taxa and across geographical settings.

## Data Accessibility

All data used in this paper are already published or publicly available.

## Author contribution statement

JNP-L and JC-B conceived the ideas presented and tested herein and contributed throughout the whole writing process.

## Acknowledgements

J.N.P-L. was supported by the University of Minnesota College of Biological Sciences’ Grand Challenges in Biology Postdoctoral Program.

## Supplementary material

**Fig. S1.**
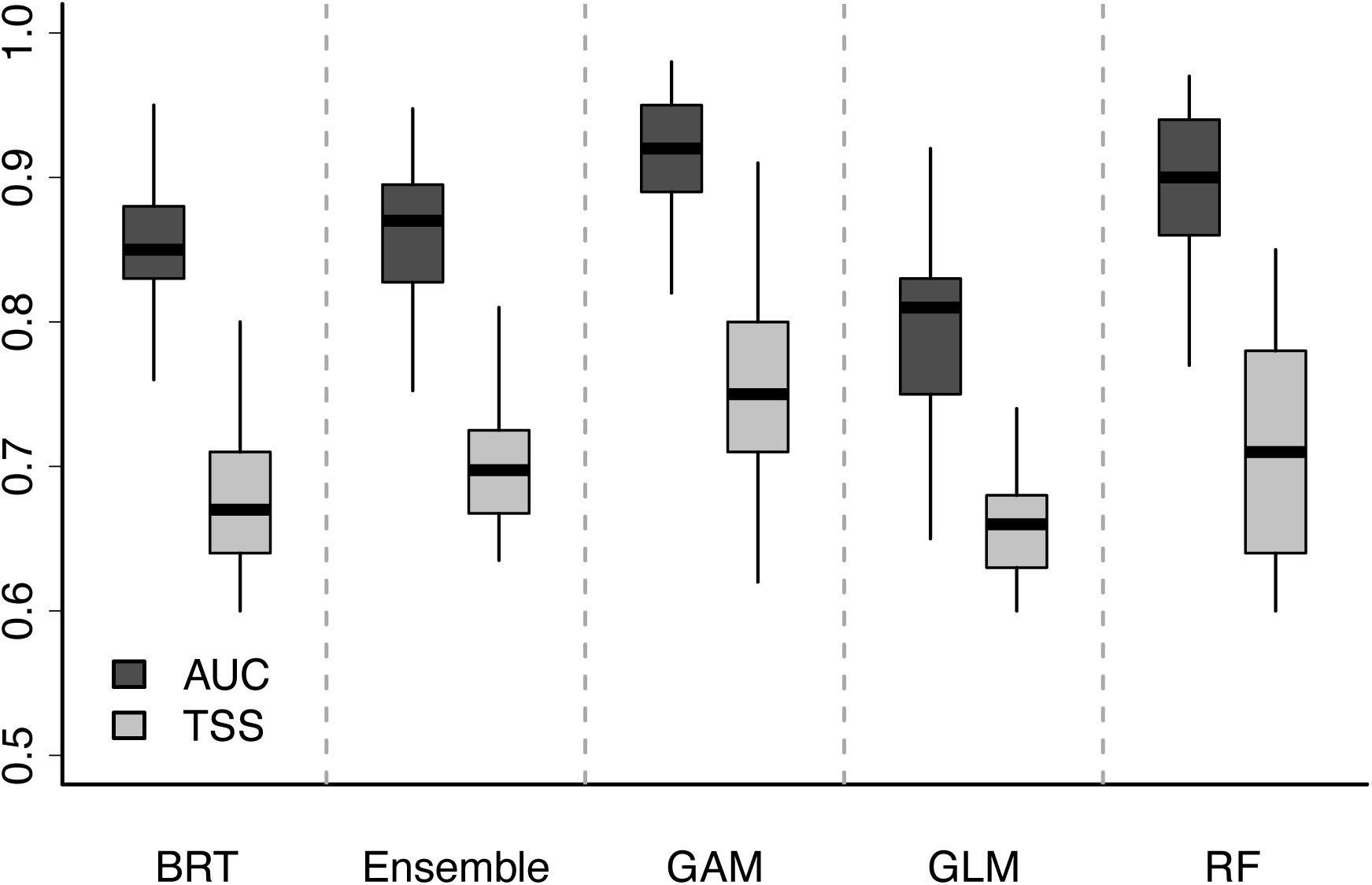
Model accuracy comparison for each algorithm and the resulting ensemble model for the 49 oak species. AUC = area under the ROC curve, TSS = true skill statistics, BRT = Boosted Regression trees, GAM = Generalized Additive Model, GLM = Generalized Linear Model, RF = Random Forest.

**Fig. S2.**
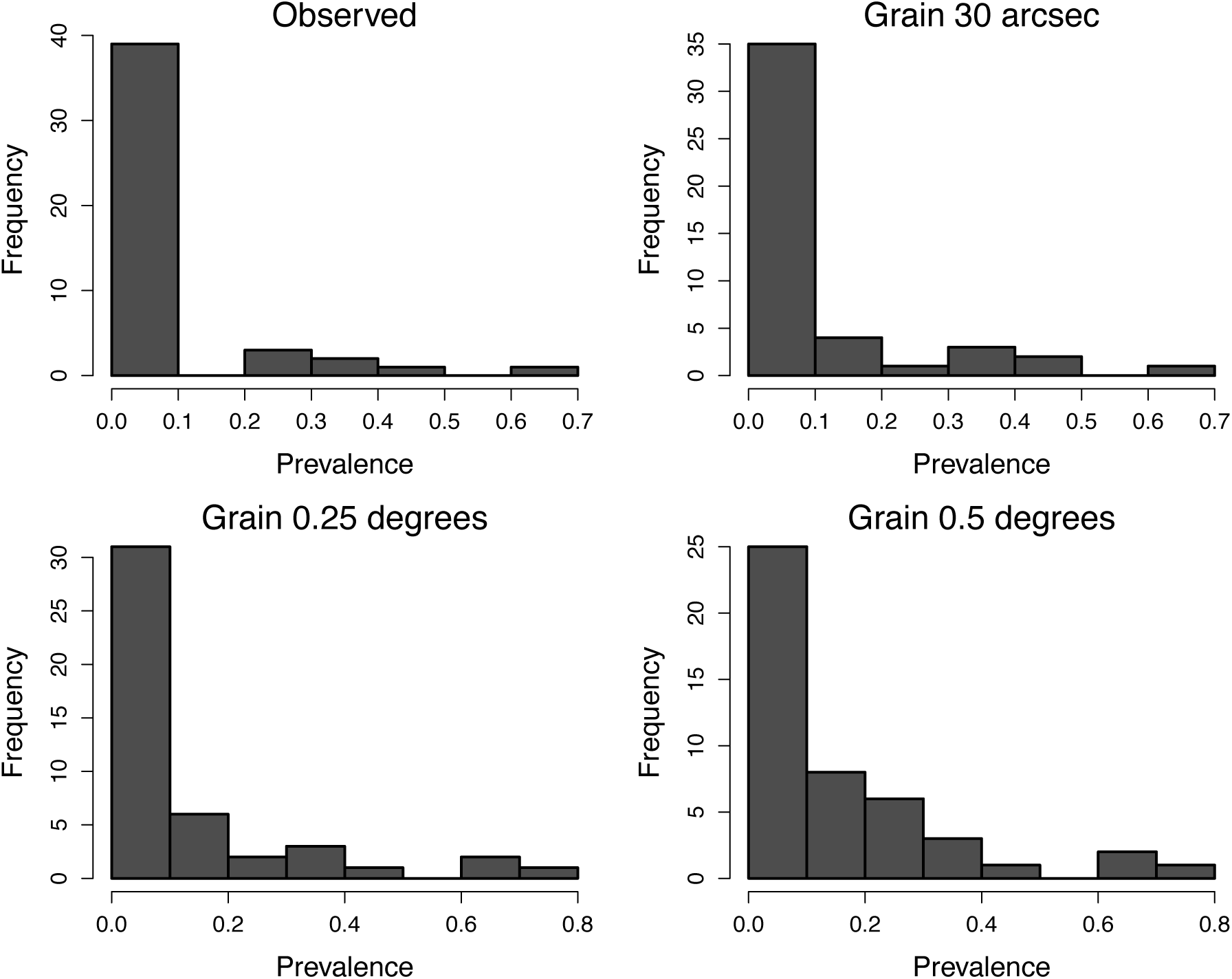
Distribution of oak tree prevalence levels for the four grain sizes estimated from the FIA dataset.

